# Peptide structural plasticity is predictable from sequence and environment

**DOI:** 10.64898/2026.08.09.743796

**Authors:** Marcelo D.T. Torres, Hanqun Cao, Cesar de la Fuente-Nunez

## Abstract

Biomolecular structure is commonly predicted from sequence as a single structural model, yet many molecules function through conformational ensembles that reorganize with their surroundings. Whether such structural responsiveness can be learned jointly from sequence and environmental context remains unresolved. Here, we use short peptides as an experimentally tractable system to test this principle. We assembled a large experimental multi-environment dataset for peptides’ secondary structure, comprising more than 1,500 peptides and more than 5,500 peptide–environment observations across aqueous, co-solvent, and membrane-mimicking conditions. We developed ApexFold, an environment-conditioned AI framework that combines sequence representations with physicochemical descriptors of the surrounding medium to predict circular-dichroism-derived fractions of ***α***-helical, ***β***-like, and unstructured conformations. In two later-collected panels excluded from model development, ApexFold captured the direction and magnitude of environment-induced structural redistribution and the peptide-specific degree of plasticity, while outperforming solvent-agnostic and composition-based baselines. Static structural references, which return a single conformation, cannot represent these condition-dependent response profiles. These results establish structural responsiveness as a learnable property of sequence and environment, extending biomolecular prediction beyond static structure toward predicting—and ultimately designing—how molecules respond to the contexts in which they function.

## 1 Introduction

Predicting molecular structure from sequence has transformed biology, but structure is not always a fixed property of sequence alone. Many flexible biomolecules populate conformational ensembles whose distributions shift as their surroundings change. Short peptides provide an experimentally tractable example: many are not best described by one dominant native structure, but instead redistribute among disordered, *α*-helical, *β*-like, and other states as solvent polarity, hydrogen-bonding capacity, membrane mimicry, and related features of chemical context change (Fig. 1a). For these molecules, the relevant question is therefore not only “what structure can this sequence adopt?” but also “for the same sequence, how does its structural ensemble reorganize when the environment changes?”[1–3].

**Fig. 1.**
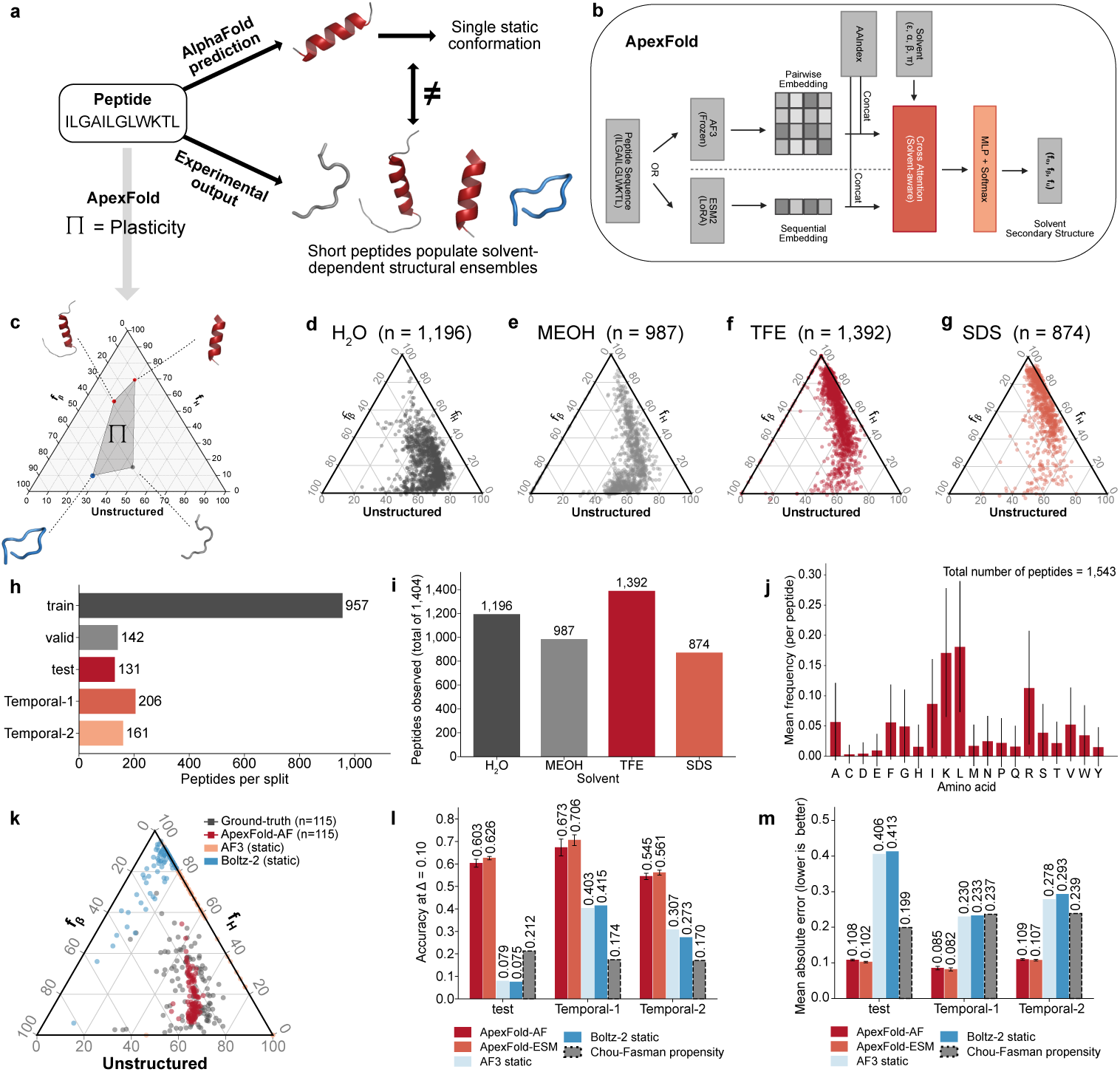
ApexFold predicts how peptide structural ensembles respond to environmental context. **a,** Short peptides can adopt a single static predicted conformation while experimentally sampling solvent-dependent structural ensembles. **b,** ApexFold architecture: AF3-frozen or ESM-2-LoRA sequence embeddings, optionally fused with AAIndex priors, conditioned on solvent descriptors (*ε, α, β, π*) by solvent-aware cross-attention and decoded to (*f_α_, f_β_, f_u_*). **c,** Per-peptide plasticity (Π): divergence among solvent-specific fraction vectors on the simplex. **d–g,** Per-solvent ternary distributions of observed (*f_α_, f_β_, f_u_*) in water, MeOH, TFE and SDS. **h,** Peptides per split. **i,** Peptides observed per solvent. **j,** Mean per-peptide amino-acid frequency (*n* = 1,543; 54 peptides were excluded from the composition calculation for containing non-canonical amino acid residues). **k,** Predicted versus observed CD fractions on the simplex: ApexFold tracks the observed cloud, whereas static references collapse to a helix-rich corner. **l,m,** Accuracy at *δ* = 0.10 and mean absolute error on `test`, Temporal-1 and Temporal-2 for ApexFold-AF and ApexFold-ESM versus static references (AF3, Boltz-2) and a Chou–Fasman baseline.

This distinction matters because molecular function can depend on context. Antimicrobial peptides, cell-penetrating peptides, and other membrane-active fragments can remain relatively disordered in water yet become helical or otherwise ordered after encountering an interface. Conversely, peptides can lose activity or selectivity when a required conformational transition is absent, excessive, or directed toward the wrong structural state. Predicting context-dependent structural responses could therefore help identify which sequences to synthesize and which conditions are most informative to text.

Circular dichroism (CD) spectroscopy provides an experimentally interpretable, population-level readout of peptide secondary structure. Here, measurements in water, methanol/water, trifluoroethanol/water, and SDS micelles allow the same peptide to be observed as its chemical surroundings change. We use these conditions as controlled experimental perturbations spanning aqueous, co-solvent, and membrane-mimicking contexts. This repeated-measurement design is crucial: it enables a model to learn structural responsiveness itself rather than a structure observed in only one condition. General-purpose structure predictors are designed primarily to recover structural models and do not directly predict CD-derived ensemble populations as a function of environment[4–6], whereas peptide-specific empirical rules are generally environment-agnostic. We address this gap with ApexFold, an environment-conditioned framework that asks whether the structural response of the same sequence can be predicted as context changes (Fig. 1b). Given a peptide sequence and a physicochemical description of the surrounding medium, ApexFold predicts the fractions of *α*-helix, *β*-like, and unstructured conformations expected in each condition and summarizes their redistribution with a structural-plasticity score (Fig. 1c). The prediction target is therefore not a single atomistic conformation, but an experimentally grounded trajectory of secondary-structure populations across environments. ApexFold is intended to complement, rather than replace, static structure prediction and experiment.

ApexFold combines learned protein representations, residue-level biophysical priors, and an explicit physicochemical representation of environment. We evaluate the framework on a multi-origin CD-derived peptide resource and, critically, on two later-collected panels that were excluded from model development(Fig. 1d-e). We ask whether the model captures not only the broad direction of structural change, but also the magnitude of redistribution and the peptide-specific degree of plasticity. More broadly, these experiments test whether structural responsiveness can be learned jointly from sequence and environmental context—a step toward molecular models that predict not only structure, but how structure changes with context.

## 2 Results

### 2.1 ApexFold predicts how structural ensembles change with environment

We reformulated peptide-structure prediction so that the output is not a single folded model, but an environment-conditioned secondary-structure ensemble. For each peptide sequence *s* and environment descriptor vector *c*, ApexFold learns a mapping to *V* = [*f_α_, f_β_, f_u_*], constrained to the probability simplex so that the three components sum to one. The same sequence can therefore receive different predicted structural populations in water, co-solvent mixtures, and membrane-mimicking media. In this formulation, context is an explicit model input rather than an unmodeled source of variation.

To quantify the magnitude of redistribution across environments, we define peptide structural plasticity (Π) as the mean pairwise Jensen–Shannon divergence among a peptide’s solvent-specific (*f_α_, f_β_, f_u_*) vectors—i.e. the Jensen–Shannon spread of its predicted or measured structural-fraction vectors across the solvents in which it is observed (Fig. 1c; see **Methods** section **Evaluation metrics** Eq. 5); a solvent-insensitive peptide has Π *≈* 0, whereas one whose ensemble redistributes strongly between conditions has large Π. This metric is bounded, symmetric, and appropriate for compositional data, making it suitable for comparing peptides whose helix, *β*-like, and unstructured fractions must change in a coupled way. When only a single environment is available, we report structural bias, *B* = 1 *− f_u_*, rather than plasticity. Plasticity was computed only for peptides with measurements in at least two solvents, because redistribution cannot be defined from a single condition.

This formulation captures the central conceptual shift of the study: the target is structural responsiveness, not simply structure in a particular solvent. ApexFold is therefore evaluated both on condition-specific structural fractions and on whether it recovers how an individual sequence changes across conditions—the direction of change, the magnitude of redistribution, and the peptide-specific degree of plasticity.

### 2.2 Repeated measurements across environments reveal structural responsiveness

We assembled a CD-derived dataset spanning natural, bioinspired, encrypted, computationally generated, and analog-optimization peptides. After curation and redundancy control, the curated development set contained 1,597 unique peptide sequences, 8 to 50 residues in length, split into training, validation and test partitions; two additional temporal panels were held out for prospective evaluation (Fig. 1h). Each peptide was measured in up to four environments (water, methanol/water, trifluoroethanol/water and SDS micelles), and the number of peptides observed in each solvent is shown in Fig. 1i. Across the dataset, all twenty amino acids were represented with substantial per-peptide variability (Fig. 1j), consistent with a compositionally diverse collection rather than a single scaffold (see **Methods** section **CD dataset construction** for details).

The key feature of this resource is its repeated measurement of the same peptides as their environments change. This design allows ApexFold to learn responsiveness itself—how a sequence redistributes its structural population across contexts—rather than associating a sequence with a structure observed in only one condition. The aqueous, co-solvent, and micellar conditions therefore serve as controlled experimental perturbations of context. Because CD deconvolution is uncertain for short peptides, we treat the task as an experimentally grounded ensemble-prediction problem and interpret performance within the resolution limits of CD-derived secondary-structure populations.

### 2.3 Environmental context enables predictions that static and environment-agnostic models do not provide

We developed two ApexFold variants that share the same environment-conditioning module and simplex-constrained prediction head. ApexFold-AF uses AlphaFold3-derived structural representations, whereas ApexFold-ESM uses embeddings from the ESM-2 protein language model. We compared these models with environment-agnostic and composition-based baselines that address the same supervised prediction problem. We also included AlphaFold3 and Boltz-2 as complementary static references (Fig. 1l,m). The two ApexFold variants showed complementary strengths. ApexFold-ESM achieved the strongest overall condition-specific performance on the later-collected panels, whereas ApexFold-AF showed particularly strong agreement for peptide-level plasticity and structural bias. Both variants substantially outperformed frozen-backbone and fully fine-tuned alternatives, indicating that pretrained protein representations can be adapted to environment-conditioned peptide prediction without relying on full-backbone fine-tuning.

### 2.4 ApexFold generalizes to unseen peptides and predicts solvent-dependent structural redistribution

We next evaluated ApexFold on two temporally held-out panels with complete four-solvent CD coverage that were excluded from model development. Sequences were partitioned along a release-time axis into three non-overlapping blocks: the earliest block was used for model development, including the internal held-out test set, while the two later blocks served as fully prospective evaluation panels, Temporal-1 (*n* = 206) and Temporal-2 (*n* = 161), ordered by release time. Because these panels postdate the development data rather than being randomly sampled from it, they evaluate generalization to peptide sequences that were excluded from model development and became available only in later collection periods.

On Temporal-1, ApexFold-ESM achieved the strongest overall performance (Acc*_δ_* = 0.706, Bias Pearson *r* = 0.509, MAE = 0.082), indicating robust transfer to peptide sequences that were not represented during model development. ApexFold-AF also generalized well (Acc*_δ_* = 0.673, Bias Pearson *r* = 0.595, MAE = 0.085) and achieved the strongest agreement in plasticity ranking (*ρ* = 0.624), suggesting that its learned structural representations capture features relevant to solvent-dependent structural redistribution beyond the training set.

A similar pattern was observed on Temporal-2. ApexFold-ESM again achieved the strongest overall predictive performance (MAE = 0.107, Acc*_δ_* = 0.561, Bias *r* = 0.644), while ApexFold-AF remained competitive in overall accuracy (MAE = 0.109, Acc*_δ_* = 0.545) and attained the highest bias correlation (Bias *r* = 0.682). Both variants substantially outperformed frozen-backbone and fully fine-tuned alternatives, indicating that peptide structural plasticity can be transferred effectively from large pretrained protein models while maintaining generalization to previously unseen peptide classes. Across both later-collected panels, ApexFold variants outperformed the composition+solvent ridge baseline and the static references in water, MeOH/water, TFE/water and SDS micelles (Fig. 2a–d). In a UMAP projection of the learned peptide representations, Temporal-1 and Temporal-2 peptides occupied the same region of representation space as the development set rather than forming isolated clusters (Fig. 2e), consistent with transfer to newly collected peptide sequences. When coloured by Pfam assignment, the same projection organized peptides into recognizable family groups (Fig. 2f). Consistent with this structure, per-family analyses showed that performance was not driven by a single dominant family: across the most populated groups, Acc*_δ_* ranged from 0.53 to 0.83 and MAE from 0.047 to 0.119, comparable to the pooled values (ALL: Acc*_δ_ ≈* 0.64–0.65, MAE *≈* 0.093; Fig. 2g,h). Finally, within-family distances were smaller than cross-family distances for sequence identity, 2-mer composition and learned ApexFold-AF/ESM profiles (Cohen’s *d* = 1.84, 2.10, 1.54 and 1.34, respectively; Fig. 2i). This separation persisted in the low-identity bin ([0.0, 0.2); Fig. 2j), indicating that the learned profiles preserve family-level organization not fully captured by pairwise sequence identity alone.

**Fig. 2.**
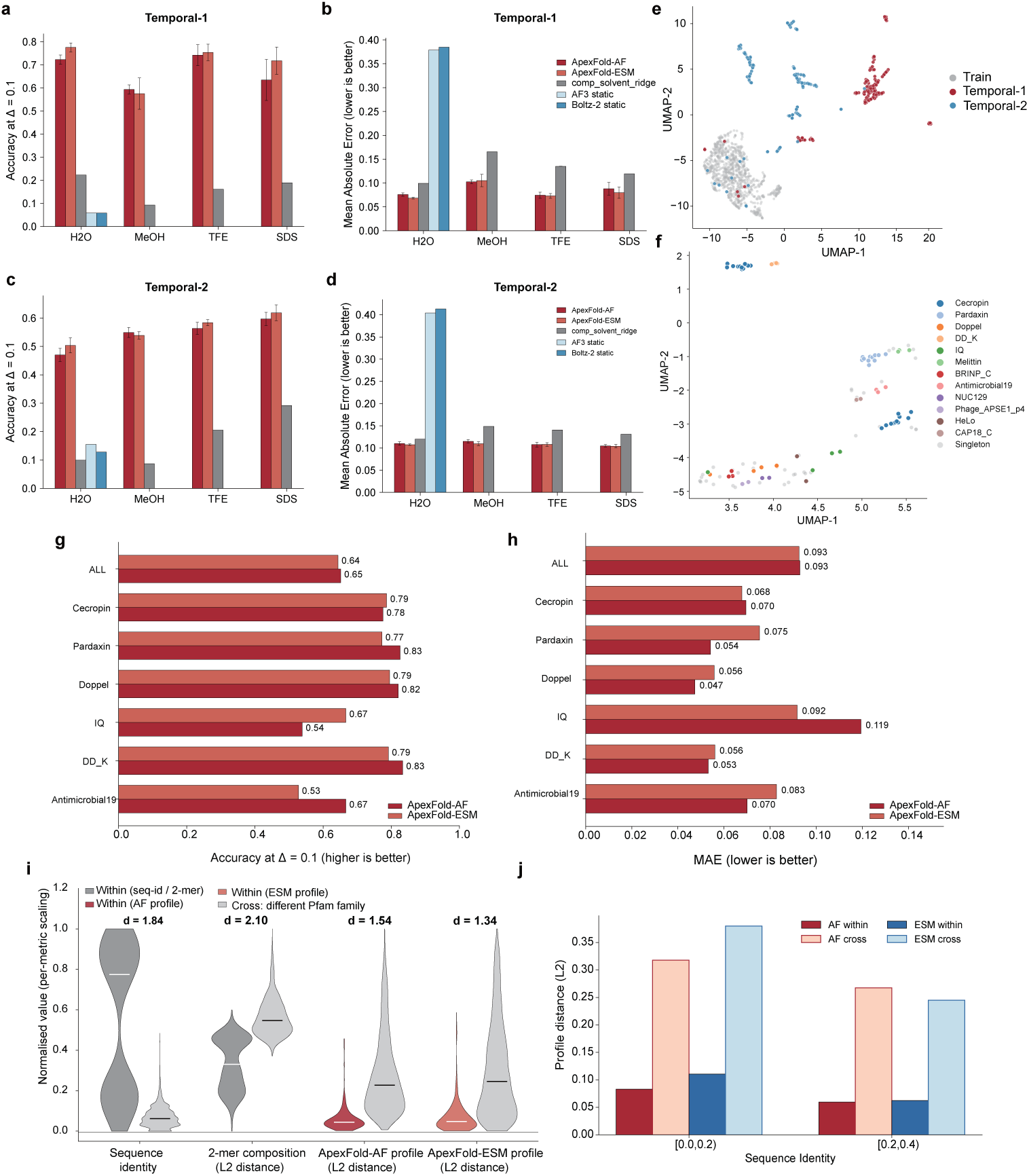
ApexFold generalizes across environments, representation space and peptide families. **a–d,** Per-solvent accuracy (Acc*_δ_*, *δ* = 0.10) and mean absolute error on **(a,b)** Temporal-1 and **(c,d)** Temporal-2 for ApexFold-AF and ApexFold-ESM versus the composition+solvent ridge and static references (AF3, Boltz-2). **e,f,** UMAP of peptide representations coloured by **(e)** split (train, Temporal-1, Temporal-2) and **(f)** Pfam family. **g,h,** Per-family accuracy and MAE for the most populated families. **i,** Within-versus cross-family distances for sequence identity, 2-mer composition and the learned ApexFold-AF/ESM profiles (Cohen’s *d* annotated).

Trend directionality was high across model families on both temporal panels (*∼* 0.95 on Temporal-2), reflecting the strong global tendency of many peptides to increase helical content in TFE or SDS relative to water. Because such global solvent trends can be captured even by simple baselines, directionality alone is not sufficient to establish peptide-specific predictive performance. We therefore interpreted directionality together with quantitative shift magnitude and plasticity metrics, which test whether the model captures how strongly individual sequences redistribute across environments.

Beyond recovering the broad direction of response, ApexFold predicted how much individual peptides changed. Predicted and observed per-residue-normalized helix shifts (Δ*f_α_*) were positively correlated across all three water-to-cosolvent transitions, with the strongest agreement observed for the more helix-promoting environments (Fig. 3a–c; Pearson *r* = 0.434/0.472/0.546 for water*→*MeOH/TFE/SDS, pooled across test, Temporal-1 and Temporal-2; *n* = 459/481/450 peptide pairs). Agreement was strongest in the more helix-promoting conditions, showing that the model captures quantitative variation in environment-induced structural redistribution rather than only the population-wide direction of change.

**Fig. 3.**
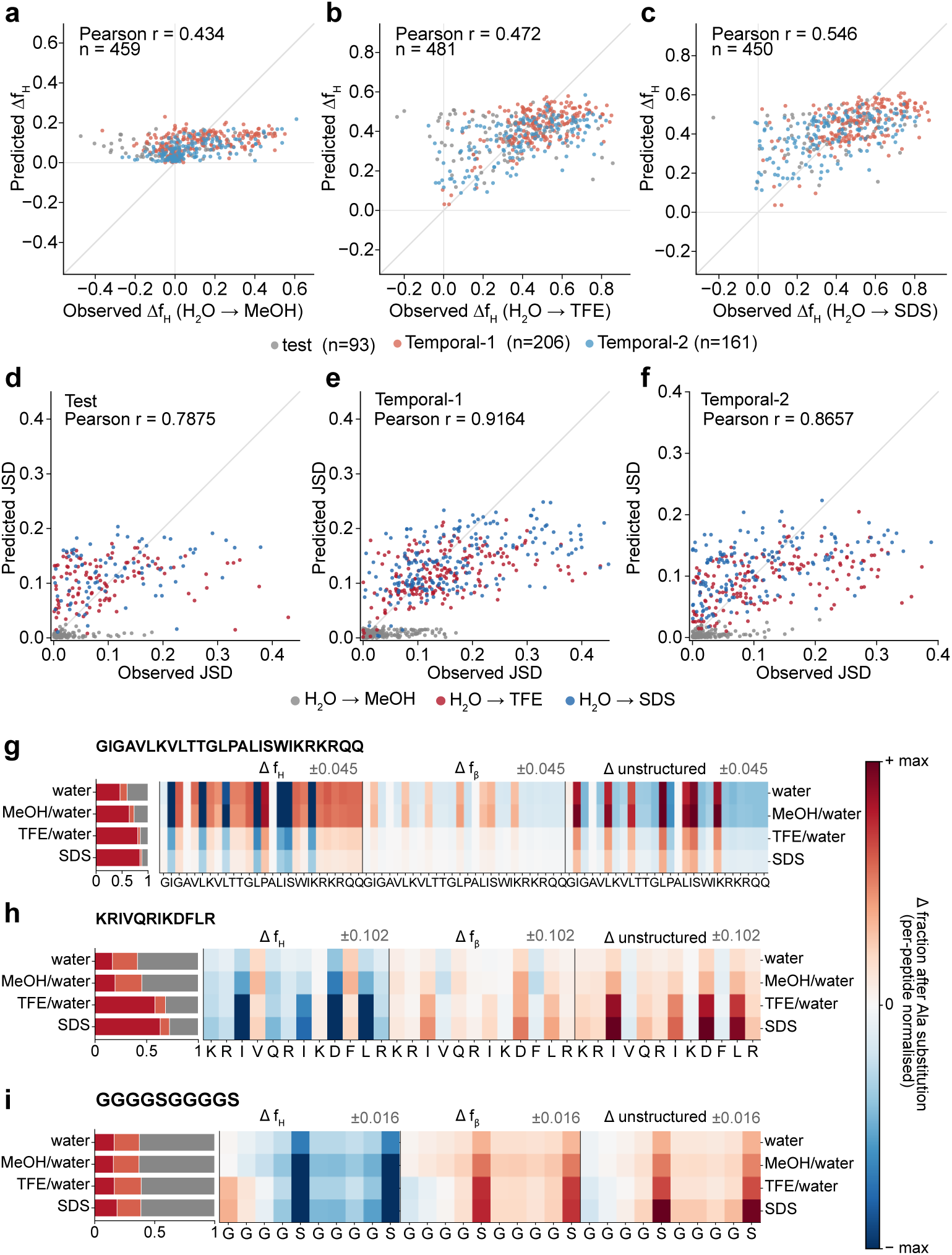
ApexFold captures the magnitude and peptide-specific degree of structural response. **a–c,** Predicted versus observed per-residue-normalized change in helix fraction (Δ*fα*) for the water*→*MeOH, water*→*TFE and water*→*SDS transitions, pooled across test, Temporal-1 and Temporal-2 (Pearson *r* = 0.434/0.472/0.546; *n* = 459/481/450 peptide pairs). **d–f,** Predicted versus observed per-peptide structural plasticity (Jensen–Shannon divergence across solvents) on test, Temporal-1 and Temporal-2 (Pearson *r* = 0.79/0.92/0.87). **g–i,** In-silico alanine-scan sensitivity maps for three example peptides (GIGAVLK_1_V_0_LTTGLPALISWIKRKRQQ, KRIVQRIKDFLR, GGGGSGGGGS): per-residue, per-solvent change in Δ*f_α_*, Δ*f_β_* and Δ*f_u_* after alanine substitution.

ApexFold also recovered the peptide-specific degree of structural plasticity (Π). Predicted and observed plasticity were strongly correlated across the internal test set and both later-collected panels (Fig. 3d–f; Pearson *r* = 0.79/0.92/0.87 for test/Temporal-1/Temporal-2). The model could therefore distinguish peptides whose structural populations remained comparatively stable from those that reorganized substantially as conditions changed. Together with the shift-magnitude analysis, this result supports the central premise that structural responsiveness can be learned from sequence and environment.

### 2.5 Sequence features associated with context-dependent structural response

To examine model behavior at the level of individual sequences, we analysed four representative peptides spanning the observed range of structural responsiveness (Fig. 4a–d). For each peptide, we overlaid the measured CD trajectory across environments with ApexFold’s condition-specific predictions and the single static conformations produced by AlphaFold3 and Boltz-2. ApexFold followed the measured movement across the structure-fraction (*f_α_, f_β_, f_u_*) simplex with low per-peptide error (MAE *≈* 0.03–0.04; e.g. IGLVIVKSISLRNPLAKFKK, ΔSS = 0.067, MAE = 0.036; IITGLSKRNKWIAKLAIKLAQK, ΔSS = 0.079, MAE = 0.041). By contrast, both static references remained fixed near the helix-rich corner regardless of environment. These examples illustrate the central limitation of static folding proxies for this task: a single environment-free conformation cannot represent peptides whose ensembles migrate between conditions.

**Fig. 4.**
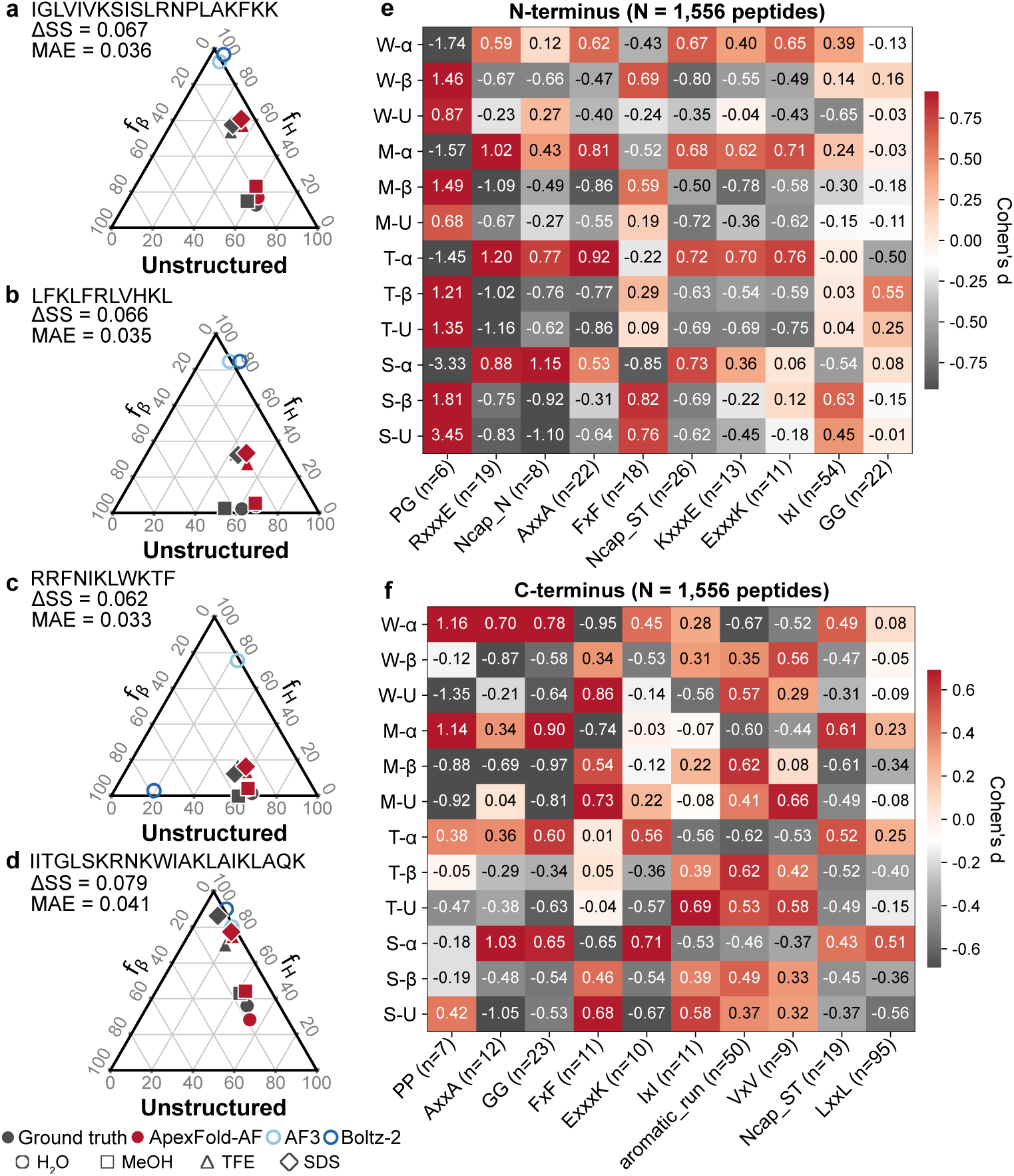
Sequence attribution identifies features associated with environment-dependent structural response. **a–d,** Four representative peptides on the (*f_α_, f_β_, f_u_*) simplex: observed CD, ApexFold predictions across solvents, and AF3 and Boltz-2 static points. ΔSS is the per-peptide secondary-structure shift magnitude across solvents; MAE is the per-peptide simplex error. ApexFold follows the measured trajectory; static references remain near the helix corner. **e,f,** Standardized associations (Cohen’s *d*) between local sequence motifs and solvent-specific structural states for **(e)** N-terminal and **(f)** C-terminal contexts. N=1,566 peptides ranging from 8 to 50 amino acid residues were computed.

We next asked which sequence features contributed to these context-dependent responses. An in-silico alanine scan perturbed each residue and measured the resulting per-solvent change in *α*-helical, *β*-like and unstructured fractions (*f_α_, f_β_, f_u_*), localizing positions to which the model assigned environment-sensitive effects (Fig. 3g–i). In parallel, we quantified associations between local sequence motifs and condition-specific structural states as standardized effect sizes (Cohen’s *d*) in N-terminal and C-terminal motif contexts (Fig. 4e–f). These associations differed systematically across environments, consistent with the model learning sequence features whose structural consequences depend of context.

These interpretability results should be viewed as mechanistic hypotheses rather than definitive causal proof. The attributions are correlational, and motif context alone did not fully explain plasticity or prediction error on the temporal splits set. The results therefore suggest that ApexFold identifies sequence features associated with environment-dependent structural response, but targeted perturbation experiments will be required to establish whether those features causally control plasticity.

### 2.6 Balanced analyses show that performance is not explained by dominant-state bias alone

Because the temporal splits set contains many *α*-rich states in TFE/water mixture and SDS micelles, raw accuracy could be inflated by predicting the dominant class. We therefore added a structure-balanced evaluation based on *α*-rich, *β*-rich, unstructured-rich, and mixed-state labels. ApexFold-AF retained the top rank after balancing, with macro Acc*_δ_* = 0.301 and dominant-state accuracy = 75.9%, while substantially exceeding global-mean and dominant-class prior baselines.

The balanced evaluation also revealed a biologically interpretable limitation. Mixed-state peptides were the hardest class, and the dominant failure mode was compression of mixed ensembles into a single dominant state. This behavior is consistent with both the three-state output representation and the uncertainty of CD deconvolution for short, flexible peptides. Rather than weakening the central conclusion, this analysis clarifies the current scope of ApexFold: it is strongest for predicting broad environment-dependentredistribution and remains less capable of resolving fine-grained mixed ensembles.

## 3 Discussion

Biomolecular prediction has largely focused on the relationship between sequence and structure, but many molecules function in environments that change. For flexible peptides, the biologically relevant object may therefore be not one structure, but an ensemble whose populations reorganize with context. By making environment an explicit model input, ApexFold shifts the prediction target from static conformation recovery toward predicting structural responsiveness—the way the same sequence redistributes its structural population as its surroundings change.

The central advance is therefore not the prediction of peptide secondary structure in particular solvents. The aqueous, co-solvent, and membrane-mimicking conditions used here are controlled perturbations through which we test a more general principle: whether responsiveness itself is learnable from sequence and environmental context. Across the internal test set and two later-collected panels, ApexFold captured the direction of structural change, the magnitude of redistribution, and the peptide-specific degree of plasticity. This matters because population-wide solvent trends can be recovered by simple baselines; the more demanding task is to predict how strongly an individual sequence responds.

This distinction also clarifies the relationship between ApexFold and modern structure-prediction systems. AlphaFold 3 and Boltz-2 were not designed to predict environment-dependent peptide ensembles or CD-derived population shifts, and our comparisons should not be interpreted as tests of their native objectives. Their static outputs instead provide a conceptual reference: one conformation cannot encode a response profile in which the populations of accessible structures change across environments. ApexFold is therefore complementary to atomistic structure prediction, adding an experimentally grounded estimate of how secondary-structure populations reorganize with context.

The immediate practical value is to make structural plasticity computationally actionable. ApexFold can help prioritize which peptides to synthesize, which conditions to test, and which candidates merit deeper structural characterization, linking high-throughput sequence discovery with lower-throughput biophysical measurement. The broader implication is that structural responsiveness itself may become a design variable. Molecular-design systems could eventually optimize not only sequence, activity, or a preferred structure, but also whether a molecule adopts the desired structural behavior in the context in which it is intended to function.

ApexFold also illustrates a strategy for combining foundation-model representations with explicit biophysical priors and experimentally defined context. Parameter-efficient adaptation, frozen-backbone representations, AlphaFold-derived pair features, and residue-level physicochemical descriptors provided complementary signals. The attribution analyses suggest that the model learns local sequence features associated with context-dependent response, but they do not establish causal mechanisms. Computational alanine scans and motif associations should therefore be used to nominate hypotheses for targeted experiments rather than interpreted as mechanistic explanations.

The current system has some limitations. CD-derived fractions are population-level consensus observables with nontrivial uncertainty, particularly for *β*-rich and mixed-state ensembles. The three-state representation groups turns, irregular structures, and rapidly interconverting coil-like states into a broad unstructured category. SDS micelles are a membrane mimic rather than a living cell membrane, and the chemical conditions studied here sample only a narrow subset of the contexts biomolecules encounter in vivo. ApexFold should therefore be interpreted as predicting broad secondary-structure redistribution across the measured environments, not an atomic-resolution folding trajectory or a universal model of biomolecular context dependence.

Short peptides provide a tractable system in which the same sequence can be measured repeatedly across controlled environments, but the broader question extends beyond peptide science: can molecular models learn how biological structure and behavior depend on context? Establishing that generality will require new datasets that perturb more physiological environments, including complex membranes, binding partners, local chemistry, and other contextual variables. The present results provide an initial proof of principle that structural responsiveness can be learned from sequence and environment at scale. A longer-term goal is therefore to predict—and ultimately design—not only the structures biomolecules can adopt, but how their ensembles reorganize in the environments in which they must function.

## 4 Methods

### 4.1 Peptide synthesis

Peptides were synthesized by solid-phase peptide synthesis (AAPPTec, Biopolymers, or in house) using standard Fmoc (9-fluorenylmethoxycarbonyl) chemistry. In-house syntheses were performed on an automated peptide synthesizer (Symphony X, Gyros Protein Technologies) using Fmoc-based SPPS on Fmoc-protected amino acid-Wang resins (100-200 mesh). N,N-dimethylformamide (DMF) was used as the primary solvent. Stock solutions consisted of 500 mmol L^-1^ Fmoc-protected amino acids in DMF, a coupling solution containing HBTU (450 mmol L^-1^) and N-methylmorpholine (900 mmol L^-1^) in DMF, and 20% piperidine in DMF for Fmoc deprotection. After chain assembly, peptides were deprotected and cleaved from the resin using trifluoroacetic acid, triisopropylsilane, dithiothreitol, and water for 2.5 h at room temperature. Peptides were precipitated with cold diethyl ether, pelleted by centrifugation, washed, dissolved in 0.1% aqueous formic acid, frozen, and lyophilized. Analytical characterization used reverse-phase HPLC and electrospray ionization mass spectrometry. Observed masses were compared with theoretical values, and quantitative analysis used integrated selected-ion-recording peak areas.

### 4.2 Circular dichroism experiments

Circular dichroism (CD) spectra were acquired to assess peptide secondary-structure content across aqueous, co-solvent and membrane-mimetic conditions[4]. Measurements were performed on a Jasco J-1500 CD spectropolarimeter at the Biological Chemistry Resource Center, University of Pennsylvania. Peptides were analysed at 50 *µ*mol l^−1^ in water, trifluoroethanol/water (3:2, v/v), methanol/water, and sodium dodecyl sulfate (SDS; 10 mmol l^−1^ in water). Spectra were collected at 25 ^◦^C using a quartz cuvette with a 1.0 mm path length over 260–190 nm, with a scan rate of 50 nm min^−1^ and a bandwidth of 0.5 nm. For each condition, three accumulations were recorded and averaged. Matching solvent baselines were acquired before the corresponding peptide measurements and subtracted from the raw spectra. A Fourier-transform filter was applied to reduce high-frequency noise and background contributions before downstream secondary-structure analysis.

### 4.3 Motivation and supervised target

ApexFold is formulated as a supervised model that predicts CD-derived secondary-structure fractions (*f_α_, f_β_, f_u_*) from a peptide sequence and a solvent descriptor vector. The supervised target is an experimentally inferred three-component vector on the simplex, not an atomistic conformation, and the model’s predictions should be interpreted at this resolution. In this work, *α*-helical, *β*-like and unstructured fractions refer to secondary-structure populations inferred from CD spectra rather than discrete atomically resolved structures. *α*-helical content denotes conformations with backbone geometry and hydrogen-bonding patterns consistent with right-handed *α*-helices, typically including *i* to *i*+4 intra-chain hydrogen bonds and CD features near 208, 222 and 190 nm. *β*-like content denotes extended backbone conformations, including *β*-strand-or *β*-like states, which in short peptides may be heterogeneous, transient or non-ideal rather than stable long-range *β*-sheets; these states are commonly associated with CD features near 215–218 and 195 nm. Unstructured content captures disordered, coil-like or rapidly interconverting conformations that do not fall into the *α*-helical or *β*-like categories and are typically associated with a minimum near 198 nm[4].

### 4.4 CD dataset construction

Peptides were compiled from in-house experimental campaigns and curated into a non-redundant sequence-level dataset. CD spectra were collected in up to four perturbation conditions: water (neat aqueous, peptide concentration 50 *µ*mol/L), methanol/water (1 : 1 v/v), TFE/water (3 : 2 v/v = 60 : 40 v/v), and SDS micelles (10 mmol/L in water). Methanol/water and TFE/water were treated as fixed solvent conditions in the supervised dataset; SDS was treated as a membrane-mimetic micellar condition rather than a physiological membrane. The Kamlet–Taft solvent descriptor table used to condition the model uses the closest tabulated mixture for methanol/water (60 : 40 v/v); this is an approximation rather than a measured property of the experimental 1 : 1 v/v mixture. The same condition definitions are used throughout the figures. The locked solvent identifier set comprises four canonical labels: water, MeOH water, TFE water, and SDS (Supplementary Table S2). Each processed CD spectrum was deconvolved into a three-component structural vector *y* = (*f_α_, f_β_, f_u_*) representing the fractions of helix, *β*-like, and unstructured structure, with *f_α_* + *f_β_* + *f_u_* = 1 and *f_k_ ≥* 0[5, 6].

The final labelled dataset comprised 1,404 measurement rows representing 1,187 unique sequences and 4,449 peptide–solvent observations across water (*n* = 1,196), methanol/water (*n* = 987), TFE/water (*n* = 1,392), SDS (*n* = 874). Solvent-specific secondary-structure distributions are shown in Extended Data Fig. 1b,c and reveal a global shift from predominantly unstructured populations in water toward increased helicity in TFE and SDS, motivating the use of solvent-conditioned prediction. The training, validation and test partitions contained 1,108/156/140 measurement rows, respectively.

### 4.5 Peptide collection and curation

The peptide collection was assembled from multiple projects over a decade and included previously reported antimicrobial peptides and analogs, some peptides that can be found in public resources such as Database of Antimicrobial Activity and Structure of peptides (DBAASP)[7], Antimicrobial Peptide Database (APD6)[8], Data Repository of Antimicrobial peptides (DRAMP)[9], UniProtKB, Arachnoserver, and ConoServer, internally curated antimicrobial peptide sets, optimization series[10–13], encrypted-peptide discovery campaigns[14–20], and computational design outputs[21–24]. This multi-origin design is emphasized in the revised manuscript because it supports the generality of the structural-redistribution question.

In the raw workbook, the collection comprised 1,597 unique amino acid sequences, with lengths ranging from 7 to 50 residues. After curation for model development and evaluation, the main dataset contained 1,187 unique sequences with measurements in up to four environments. The sequence-length distribution per split, and the distributions of key sequence-level physicochemical descriptors of the curated dataset—net charge, hydropathy (GRAVY), isoelectric point, molecular weight, aromaticity, instability index and Boman index—are shown in Extended Data Fig. 1a,d–k, and confirm that the atlas spans a broad, predominantly cationic and moderately hydrophobic peptide space rather than a single scaffold. Redundancy was controlled using CD-HIT clustering at 70% sequence identity, with clusters allocated to train, validation, and test splits.

### 4.6 Model formulation

ApexFold learns a mapping *f_θ_*(*s, c*) *→ V*, where *s* is a peptide sequence, *c* is a continuous solvent descriptor vector, and *V* = [*f_α_, f_β_, f_u_*] is a secondary-structure fraction vector constrained to the simplex. The output is therefore directly comparable to CD-derived ensemble fractions.

Solvents were represented using dielectric constant and Kamlet-Taft solvatochromic parameters. For SDS micelles, these values should be interpreted as effective interfacial descriptors that capture the average CD-observable outcome of peptide-micelle interaction, not as homogeneous bulk solvent parameters.

### 4.7 Plasticity metric

Structural plasticity was quantified as the mean pairwise Jensen-Shannon divergence among the solvent-specific secondary-structure fraction vectors associated with each peptide. Plasticity therefore measures the extent to which a peptide redistributes its structural population across environments, with low values indicating similar structural compositions across solvents and high values indicating substantial solvent-dependent redistribution. Peptides measured in fewer than two solvents were excluded because redistribution cannot be defined from a single condition (Fig. 1c). Accordingly, Π is used as a peptide-level summary of structural responsiveness across the measured environmental panel. It quantifies redistribution of the CD-derived population vector and should not be interpreted as an atomistic measure of conformational entropy.

### 4.8 Model architecture and training

ApexFold predicts the four-solvent secondary-structure simplex of a peptide in a single forward pass. The architecture decomposes into (i) a multi-source sequence encoder, (ii) a solvent-aware cross-attention conditioning module, and (iii) a simplex-constrained prediction head. Training combines a missingness-aware reconstruction loss with an auxiliary plasticity-matching term (architecture overview in Fig. 1b).

#### 4.8.1 Sequence representation

Each peptide is encoded as a concatenation of three complementary residue-level views, providing both learned semantic context and inductive physicochemical priors. First, ESM-2 embeddings: we extract per-residue hidden states from a frozen ESM-2 backbone (esm2 t30 150M UR50D, 30 transformer layers, hidden dimension 640) at the final layer, giving an *L* × 640 matrix that supplies evolutionary and contextual semantics learned from UR50; the effect of the extraction layer and of the backbone size on accuracy is characterised in Extended Data Fig. 2c–e. Second, AF3 representations: for each peptide we run AlphaFold-3 in static (no-MSA) mode and harvest the last-layer single representation *s ∈* ℝ*^L^*^×384^ and pair representation *z ∈* ℝ^*L*×*L*×128^; the single track contributes residue-level fold preferences, whereas the pair track encodes positional co-dependencies that a secondary-structure-fraction predictor can exploit (helix-stabilising *i, i* + 4 contacts and *β*-bridge *i, i ± n* motifs). Consistent with this interpretation, the AF3 pair representation is dominated by short-range, near-diagonal interactions (Extended Data Fig. 2i), and progressively masking it reduces prediction accuracy (Extended Data Fig. 2h). Third, biophysical priors: we append a 21-dimensional residue-level descriptor derived from AAIndex1, comprising hydrophobicity (Kyte–Doolittle), helix, *β*-like and coil propensities (Chou–Fasman), volume, polarity, charge, isoelectric point, and *α*-helix and *β*-strand free-energy contributions. These hand-crafted features supply an inductive bias toward classical sequence-to-structure rules and stabilise learning on small datasets; the complementary contribution of each of the three feature classes, individually and in combination, is quantified in Extended Data Fig. 2a,b. The three views are projected to a common width *d*_model_ = 256 through learned linear maps and summed token-wise to form a unified residue embedding **h** *∈* ℝ^*L*×*d*_model_^.

#### 4.8.2 Solvent-aware cross-attention conditioning

Solvent identity (water, MeOH/water, TFE/water or SDS) is represented as a learned 64-dimensional embedding **e***_s_* concatenated with three scalar physicochemical descriptors (dielectric constant *ε*, surface tension *γ*, and a fluorinated/anionic indicator), then passed through a two-layer MLP to produce a solvent query vector **q***_s_ ∈* ℝ^*d*_model_^. A two-layer cross-attention module conditions peptide tokens on solvent context:

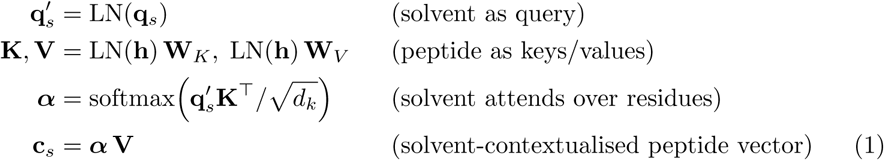

The output **c***_s_ ∈* ℝ^*d*_model_^ is a peptide summary as seen through that solvent. This direction (solvent queries peptide, rather than peptide queries solvent) ensures that one solvent forward pass yields one global readout, matching the experimental observable of a single CD spectrum per peptide-solvent pair and avoiding per-residue solvent ambiguity that the labels cannot supervise. To produce all four solvent predictions in a single forward pass, we stack the four solvent queries into a 4 *× d*_model_ batch and decode them in parallel.

#### 4.8.3 Simplex-constrained prediction head

Each solvent-contextualised vector **c***_s_* is mapped through a three-layer MLP (GELU activation, dropout 0.2) to logits **z***_s_ ∈* ℝ^3^, which are passed through a softmax to enforce the probability-simplex constraint *f_α_* + *f_β_* + *f_U_* = 1 with *f ∈* [0, 1]^3^. We deliberately avoid an unconstrained sigmoid with post-hoc renormalisation, because the latter weakens the gradient signal at the boundary of the simplex, where helix-dominant and disorder-dominant peptides live. The output is the 4 × 3 matrix **F̂** of predicted fractions across *{*water, MeOH/water, TFE/water, SDS*} × {α, β, U }*.

#### 4.8.4 Training objective

Let **F** be the observed fractions, **M** *∈ {*0, 1*}*^4×3^ the per-cell observation mask, and **F̂** the model prediction. The total loss combines three terms:

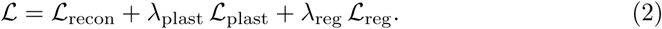

**Masked reconstruction..**

(*L*_recon_). We use a smooth-*L*_1_ (Huber, *β* = 0.05) loss on the simplex, applied only to observed cells and normalized by the mask sum so that under-supervised peptides are not down-weighted:

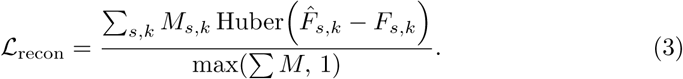

The Huber loss is preferred over MSE because CD-derived fractions carry heavy-tailed noise from baseline drift and basis-set choice (the inter-method spread across BestSel, CDSSTR, CONTINLL and SELCON3); an *ℓ*_2_ loss over-weights these tails and degrades calibration.

**Plasticity-matching auxiliary..**

(*L*_plast_). To encourage the model to learn conformational shifts rather than only per-cell calibration, we add a divergence-matching loss over solvent pairs. For each ordered solvent pair (*s, s*^′^) with both members observed, we compute the Jensen–Shannon divergence of the predicted and observed fraction triples and penalise their squared difference:

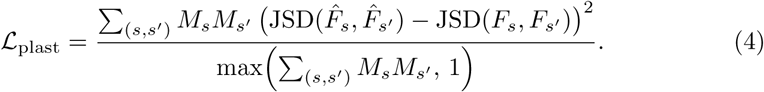

This term directly supervises the plasticity ranking (*ρ*) reported in the main text and is important for predicting AMP-like peptides whose biological function depends on a water-to-SDS helical switch rather than on absolute fractions.

**Logit regularisation..**

(*L*_reg_). A small *ℓ*_2_ penalty on the pre-softmax logits (*λ*_reg_ = 10^−4^) prevents the simplex head from saturating early in training, where the softmax gradient vanishes near the vertices.

We set *λ*_plast_ = 0.5 by validation-set grid search; values in [0.3, 0.8] performed comparably.

#### 4.8.5 Optimization

We use AdamW with learning rate 3×10^−4^ (linear warmup over 500 steps, cosine decay to 3 × 10^−5^), weight decay 10^−2^, batch size 32, and gradient clipping at 1.0. Training runs for 200 epochs with early stopping on validation MAE (patience 20). With mixed-precision (bfloat16) on a single A100 GPU, full training of one seed completes in approximately 1.5 h for ApexFold-AF and approximately 4 h for ApexFold-ESM. Three seeds (42, 123 and 456) are trained per configuration and metrics are reported as mean *±* s.d.

#### 4.8.6 Handling label heterogeneity

Ground-truth fractions are aggregated across the four CD deconvolution methods (BestSel, CDSSTR, CONTINLL and SELCON3) by per-cell mean. Cells where all four methods fail (typically SDS *β*-content under high-noise spectra) are masked out of *L*_recon_ and *L*_plast_ through **M**. An ablation trained on BestSel-only labels is reported in the Supplementary Information as a label-source sensitivity check.

### 4.9 Train, validation, test and temporal splits

Model-development sequences were divided into training, validation and held-out test partitions at the peptide level. Sequence variants and near-duplicates were grouped before splitting using CD-HIT clustering at 70% sequence identity; cluster membership was used to enforce zero cluster overlap between training, validation, test and temporal splits partitions[25]. Sequence-identity-level leakage diagnostics for the temporal splits are reported in Supplementary Table S5: the mean nearest-train identity is 0.42 for Temporal-1 and 0.37 for Temporal-2, and only 1.9% and 0.6% of Temporal-1 and Temporal-2 peptides have any nearest-train neighbor at *≥* 0.7 identity. Pfam family labels were assigned computationally (see “Pfam family assignment and per-family analysis”) and are used only for descriptive, group-conditional analyses; campaign and biosynthetic-source metadata are encoded in peptide names by convention rather than as structured columns, and no field was used to define held-out groups. We therefore report sequence-identity-level leakage diagnostics and group-conditional analyses, and do not claim true leave-group-out validation in this work. We therefore report only sequence-identity-level leakage diagnostics and group-conditional analyses; we do not claim true leave-group-out validation in this work. Independent temporal splits panels were excluded from model development and used only for external evaluation: an antimicrobial peptide panel (Temporal-1) and an encrypted peptide panel (Temporal-2). Sequence-length distributions were comparable across the training, validation, test and temporal panels (Extended Data Fig. 1a), indicating that the temporal split does not confound generalization with a shift in peptide length. Final primary denominators after the deduplicated evaluation protocol are test *n* = 131, Temporal-1 *n* = 206 and Temporal-2 *n* = 161 (see “Primary deduplicated evaluation protocol” above).

### 4.10 Pfam family assignment and per-family analysis

To characterise how peptides are organised in representation space, we assigned each sequence to a protein family by scanning it against the Pfam-A database with HMMER (hmmscan) and retaining the top-scoring hit below an E-value of 1.0; sequences without a significant hit were labelled Singleton. These family labels were used only for the descriptive analyses in Fig. 2f–j and were never used to define the training, validation, test or temporal splits; per-family performance and within-versus cross-family distances are therefore reported as group-conditional descriptive analyses rather than as leave-one-family-out generalisation. Per-family accuracy and MAE (Fig. 2g,h) were computed for the most populated families. For the within-versus cross-family analysis (Fig. 2i,j), each peptide pair was scored by sequence-identity distance, the *L*_2_ distance between 2-mer composition vectors, and the *L*_2_ distance between the learned ApexFold-AF and ApexFold-ESM peptide profiles, each defined as the concatenation of the predicted (*f_α_, f_β_, f_u_*) vectors across the four solvents. pairs were grouped into within-family and cross-family sets, summarised as standardised effect sizes (Cohen’s *d*), and additionally stratified by sequence-identity bin to test whether family separation in the learned profiles persists when sequence similarity is held low.

### 4.11 Solvent representation

Each solvent condition was represented by a combination of categorical identity and physicochemical descriptors capturing solvent polarity, hydrogen-bond donor/acceptor parameters and an environment-class indicator, following solvatochromic descriptor concepts and micellar polarity measurements where applicable[26–28]. For interpolation analyses, descriptors were linearly interpolated between endpoint environments and passed through the trained model without retraining; these interpolated predictions are model-based hypotheses, not measurements, and are not validated against intermediate experimental compositions.

### 4.12 ApexFold architecture

ApexFold consists of a sequence encoder, a solvent encoder and a prediction head. The sequence encoder uses embeddings from a pretrained protein language model (ESM-2 backbone, with backbone size and adaptation strategy reported in Supplementary Tables S2 and S15)[29]. The solvent encoder maps the solvent descriptor vector to a learned representation. Sequence and solvent representations are fused by a lightweight cross-attention prediction module to produce three logits corresponding to helix, *β*-like, and unstructured fractions. A softmax converts logits to normalized fractions on the simplex. Parameter-efficient variants use low-rank adaptation of selected language-model layers while keeping most pretrained parameters frozen[30]; the effect of the low-rank-adaptation rank on accuracy across panels is shown in Extended Data Fig. 2f.

### 4.13 Training objective and model selection

Models were trained to minimize the discrepancy between predicted and CD-derived structural fractions. The primary loss combined component-wise regression error with the simplex constraint induced by the softmax output. Validation performance was monitored using mean absolute error, component-wise accuracy at fixed tolerance and structural-bias correlation. Model selection prioritized a combination of validation performance and temporal splits stability (selection criterion: best mean of test and temporal splits Acc*_δ_* at *δ* = 0.10), rather than training loss alone. Hyperparameters included learning rate, low-rank adaptation rank, dropout, batch size and the relative weighting of sequence and solvent features.

### 4.14 Evaluation metrics

Let *ŷ_i,s_* and *y_i,s_* denote the predicted and measured (*f_α_, f_β_, f_u_*) vectors for peptide *i* in solvent *s*. Component-wise mean absolute error was computed across the three structural fractions. Accuracy at tolerance *δ*, denoted Acc*_δ_*, is defined as the fraction of structural components for which *|ŷ_i,s,k_ − y_i,s,k_| ≤ δ*; we report Acc*_δ_* at *δ ∈ {*0.05, 0.10, 0.15, 0.20, 0.25*}* (Supplementary Table S1) to avoid threshold-specific conclusions, alongside continuous MAE/RMSE. We interpret Acc*_δ_* as a tolerance-based agreement measure under CD deconvolution noise rather than as a theoretical ceiling for a perfect predictor; the inter-algorithm CD-deconvolution noise ceiling is included in Supplementary Table S1 for context. Structural bias is summarized as *B* = 1 *− f_u_* (the ordered, i.e. non-unstructured, fraction), and Pearson correlations between predicted and measured *B* are computed across sequence–solvent observations.

The operational plasticity score for a peptide *i* is defined as the Jensen–Shannon divergence among its solvent-specific (*f_α_, f_β_, f_u_*) vectors:

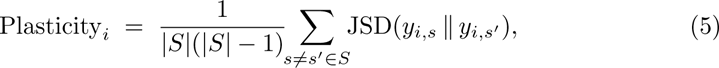

where *S* is the set of solvents in which peptide *i* was measured. Plasticity is a divergence on the simplex, not a measure of atomistic conformational entropy. Directional consistency is evaluated by comparing the sign of predicted and measured changes in each structural fraction between solvent pairs. Peptides with *|S| <* 2 were excluded from plasticity analyses.

To avoid attributing trivially recoverable solvent effects to ApexFold, response-level metrics are reported alongside three reference baselines (Supplementary Table S3): (i) a solvent-only baseline that predicts the per-solvent dataset mean of (*f_α_, f_β_, f_u_*) regardless of sequence; (ii) a per-(source,target) mean-shift (“global-transition”) baseline that adds the dataset-average Δ(*f_α_, f_β_, f_u_*) between two solvents to the aqueous prediction; and (iii) a composition-plus-solvent ridge regression (comp solvent ridge) trained on 20-D amino-acid composition with per-solvent weights. We additionally report a sequence-shuffled control that retrains a sequence model on a permuted sequence-to-label mapping (seq shuffled ESM-30+AA); on the primary deduplicated test set this control attains MAE = 0.156 versus 0.149 for the unshuffled ESM-30+AA model, and on Temporal-1 MAE = 0.222 versus 0.175, confirming that the sequence-to-fraction mapping is non-trivial. Static-structure proxies (Chou–Fasman propensity scoring; Boltz-2 single-structure outputs decomposed into secondary-structure fractions) are reported as static-structure references rather than competitors and consistently underperform the trivial solvent-only mean by a factor of two to three on simplex MAE (Supplementary Table S3)[31, 32]; their inclusion serves to test whether a single environment-free conformation can substitute for solvent-conditioned CD fractions, not to evaluate AlphaFold 3 or Boltz-2 on their native folding tasks.

All Acc*_δ_*, MAE, RMSE and dominant-state accuracy values reported in the main figures and tables are accompanied by 95% confidence intervals from a peptide-level non-parametric bootstrap (10,000 resamples, default seed 20,260,429): peptide identifiers are resampled with replacement and all condition rows for a peptide travel together, which preserves the clustered design effect (peptides measured in multiple solvents are correlated). Acc*_δ_* is reported at *δ ∈ {*0.05, 0.075, 0.10, 0.125, 0.15, 0.20*}* (Supplementary Table S6) so that no single threshold drives the conclusion. Inter-deconvolution-algorithm uncertainty on the CD-derived fractions sets a floor on what tolerance-based agreement metrics can resolve: scoring all peptide × solvent × component tuples across BestSel, CDSSTR, CONTINLL and SELCON3 yields a median between-algorithm SD of 0.094, with 47% of tuples showing SD *>* 0.10 and 11% showing at least one out-of-simplex algorithm value (negative component or sum *>* 1.05); see Supplementary Table S7 and Extended Data Fig. S8. We use *δ* = 0.10 as the primary operating point for Acc*_δ_* to remain well above this empirical floor.

For trend metrics, for each ordered solvent pair (src, dst) we computed per-peptide Δ = (*f*_dst_ *− f*_src_) and reported (a) directional agreement (sign(Δ̂) = sign(Δ) at threshold *|*Δ*| >* 10^−6^) and (b) the Pearson correlation *r*(Δ) between predicted and measured shift magnitudes. Both metrics are reported alongside the same three reference baselines defined above. Solvent-only directional-agreement values of 0.78–0.82 across panels define a trivial-baseline ceiling against which ApexFold’s directional agreement is not informative; we therefore use *r*(Δ) as the primary discriminating metric (Supplementary Table S4).

### 4.15 Static-proxy baseline interpretation

Static structure-prediction baselines were evaluated as static proxies rather than as direct competitors trained for the same supervised target. AlphaFold 3 and Boltz-derived single-structure outputs do not provide solvent-conditioned CD fractions; we converted them into coarse secondary-structure summaries only to test whether a single environment-free conformation can approximate measured CD ensembles. The conclusion from this comparison is that static aqueous summaries are not sufficient substitutes for solvent-conditioned CD-fraction prediction, not that AlphaFold 3 or Boltz-2 fail at the tasks for which they were designed.

### 4.16 Feature ablations

We ablated the contribution of each sequence-feature class to quantify their complementary value (Extended Data Fig. 2a,b). AAIndex-derived residue features improved generalization for ESM-family models, especially on OOD sequences, where they acted as a regularizing biophysical prior. For AF3-derived features, the benefit depended on representation type: AAIndex improved AF3-s and improved OOD performance of AF3-z, but it could reduce in-distribution AF3-z accuracy, suggesting that the pair representation already encodes part of the same local structural signal (Extended Data Fig. 2h,i). These ablations indicate that peptide structural plasticity is not captured by a single feature class: language-model embeddings provide broad sequence context, explicit biophysical priors stabilize learning in small-data regimes, and AF3-derived pair features encode local geometry that is especially useful for ranking plasticity. The best practical configuration therefore depends on whether the priority is scalability, OOD generalization, or plasticity ranking. Beyond feature class, we separately examined the ESM-2 extraction layer (Extended Data Fig. 2c), the ESM-2 backbone size (Extended Data Fig. 2d,e), the low-rank-adaptation rank (Extended Data Fig. 2f) and the training-loss aggregation scheme (Extended Data Fig. 2g), as well as the internal structure of the AF3 pair representation (Extended Data Fig. 2h,i), to fix the final ApexFold-AF and ApexFold-ESM configurations. In the loss-aggregation comparison, “normalized” denotes mean-aggregating the reconstruction term *L*_recon_ over observed cells while leaving the auxiliary plasticity term *L*_plast_ unchanged; because this helped the smaller ESM-2 backbone but not the larger one, we retained the unnormalized loss as the default.

### 4.17 Quality control and supplementary analyses

Supplementary analyses included dataset and split ablations, amino-acid-index correlations, solvent-shift analyses, simplex visualizations, leakage diagnostics, sigmoid fits to titration-like curves, and correlation matrices among CD-derived quantities[33]. These analyses verify that the model is not driven solely by composition, peptide length or dataset imbalance, and they identify sequence features associated with solvent-induced redistribution.

## Extended Data and Supplementary Figures

## Data availability

All data needed to reproduce the findings of this study are openly available. The circular-dichroism measurements, the deconvolved per-peptide secondary-structure fractions (*f_α_, f_β_, f_u_*) for all four solvents, the sequence-identity-based train/validation/test and Temporal-1/Temporal-2 split assignments, the sequence-clustering files, and the ApexFold predictions for the held-out test and temporal panels have been deposited in a permanent repository https://zenodo.org/records/20616716. Source data for every main-text and Supplementary figure are provided with the paper. The AlphaFold-3 and Boltz-2 structural embeddings used as model inputs are large binary files that can be regenerated from the released sequences using the published AlphaFold-3 and Boltz-2 inference pipelines[32, 34], and the ESM-2 embeddings can be regenerated from the same sequences with the ESM-2 model[29].

## Code availability

The ApexFold training and inference code, the trained model checkpoints, and the analysis scripts used to reproduce all results, figures and Supplementary tables are available at https://gitlab.com/machine-biology-group-public/apexfold and archived with a permanent identifier https://zenodo.org/records/20616716. The repository includes documentation and the commands needed to reproduce every reported metric.

## Acknowledgements

Cesar de la Fuente-Nunez holds a Presidential Professorship at the University of Pennsylvania. Research in this publication was supported by the National Institute of General Medical Sciences of the National Institutes of Health under award number R35GM138201 and the Defense Threat Reduction Agency (DTRA; HDTRA1-21-1-0014). We thank members of the Machine Biology Group for discussions.

## Author contributions

M.D.T.T., H.C. and C.d.l.F.-N. conceived the study. M.D.T.T. performed all the circular dichroism experiments. M.D.T.T. and H.C. assembled datasets, developed models, performed analyses and prepared figures. C.d.l.F.-N. supervised the project. All authors contributed to writing and approved the manuscript.

---

**Extended Data Fig. 1.**
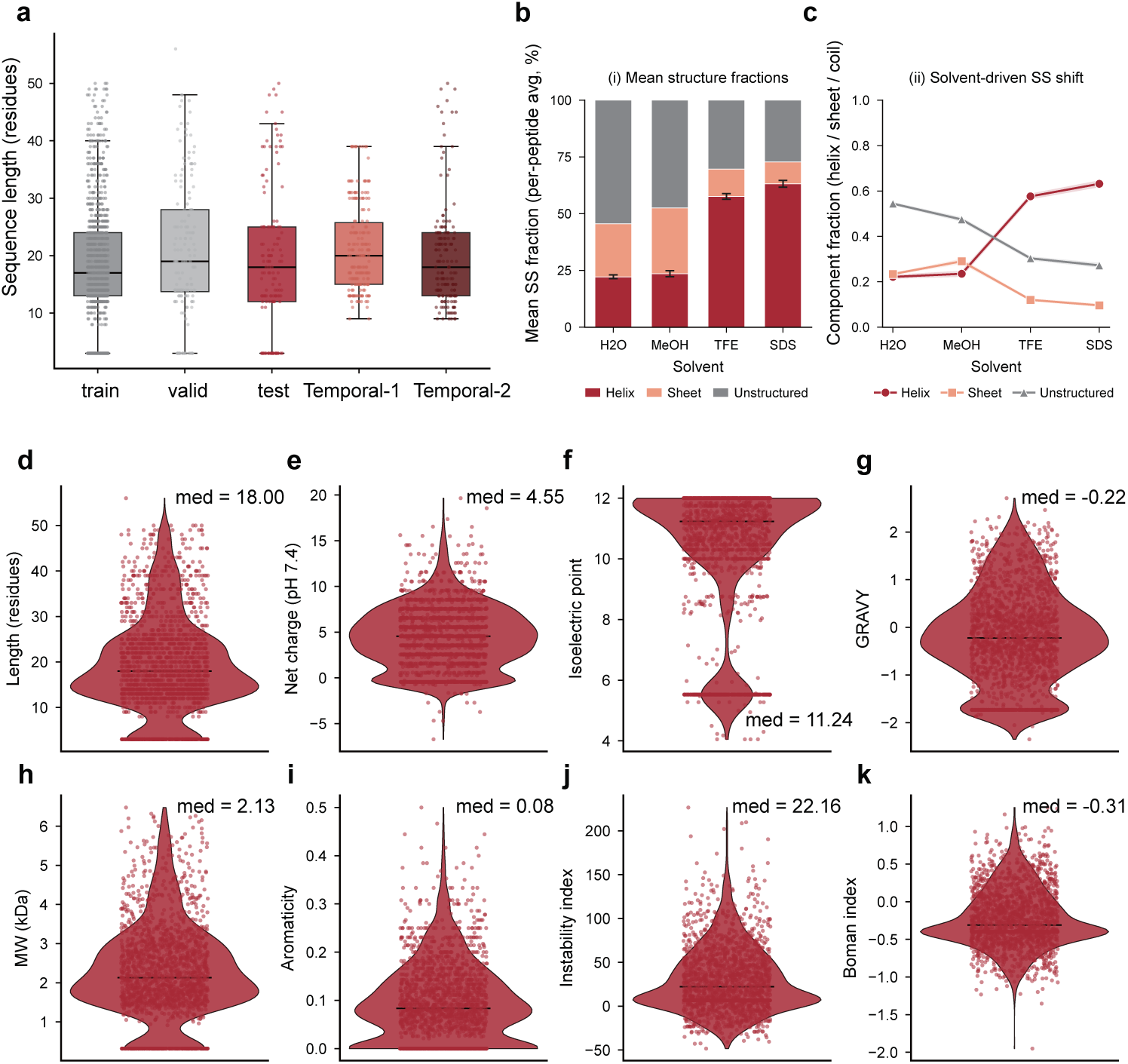
Composition and physicochemical characterization of the multi-solvent CD atlas. **a,** Sequence-length distribution (residues) for the training, validation, test, Temporal-1 and Temporal-2 splits (box: interquartile range and median), showing that peptide length is comparable across the development and prospective panels. **b,** Mean per-peptide secondary-structure composition (helix, *β*-like, unstructured; %) in each solvent (water, MeOH/water, TFE/water, SDS), summarising the dataset-level structural bias of every condition. **c,** Solvent-driven secondary-structure shift: mean helix, *β*-like and unstructured fractions across water *→* MeOH *→* TFE *→* SDS, showing the systematic increase in helix and decrease in unstructured content as the medium becomes more helix-inducing or membrane-like. **d–k,** Distributions of sequence-level physicochemical descriptors over the curated dataset (median annotated): **(d)** length, **(e)** net charge at pH 7.4, **(f)** isoelectric point, **(g)** GRAVY hydropathy, **(h)** molecular weight, **(i)** aromaticity, **(j)** instability index and **(k)** Boman index. Together these panels show that the atlas spans a broad, predominantly cationic and moderately hydrophobic peptide space rather than a single scaffold.

**Extended Data Fig. 2.**
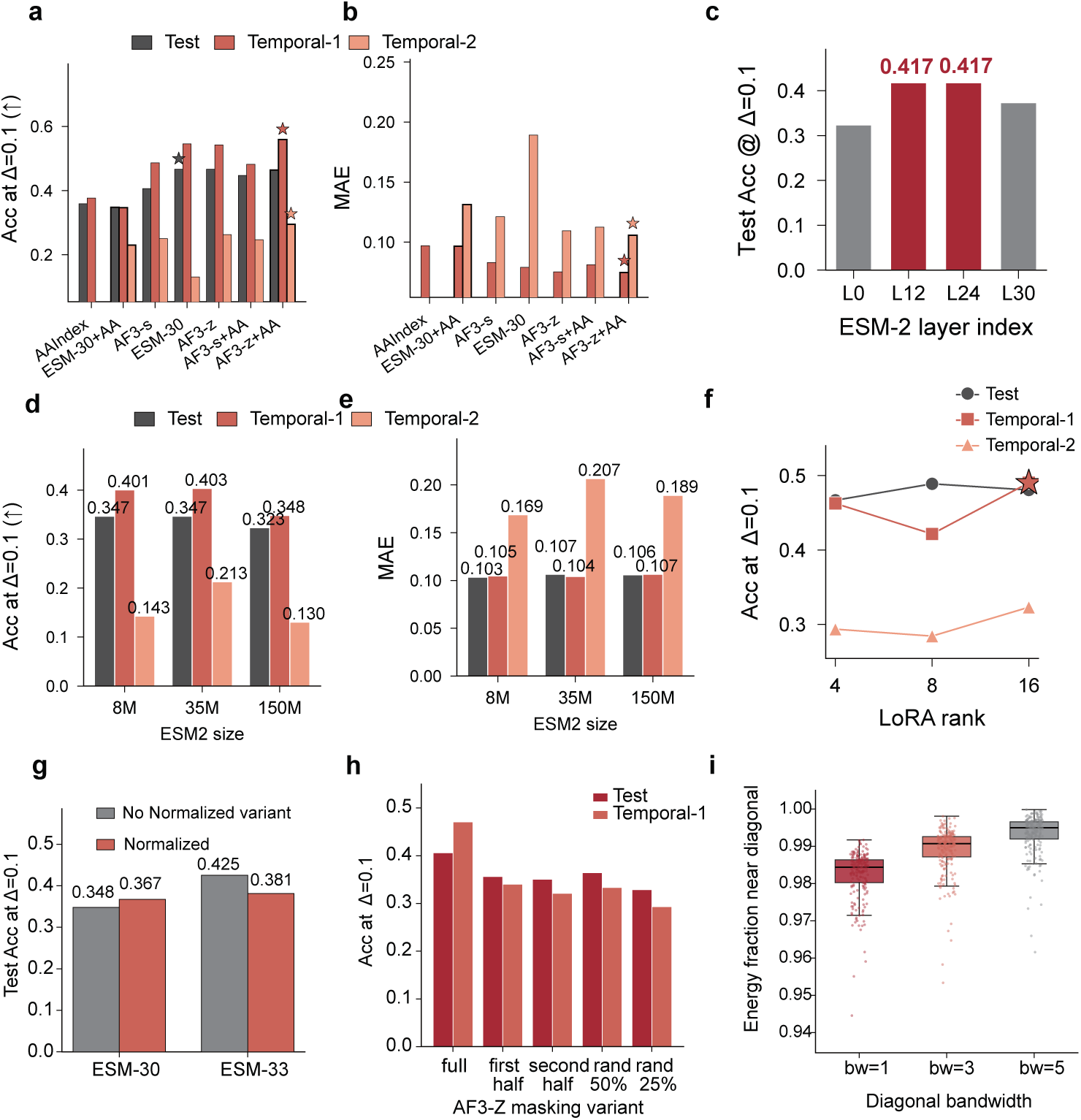
Feature-encoding and architecture ablations for ApexFold. **a,b,** Accuracy at *δ* = 0.10 **(a)** and mean absolute error **(b)** on the `test`, Temporal-1 and Temporal-2 panels for individual and combined sequence-feature classes: AAIndex priors alone; ESM-2 (layer-30) embeddings with and without AAIndex (ESM-30, ESM-30+AA); and AF3 single- and pair-track features with and without AAIndex (AF3-s, AF3-s+AA, AF3-z, AF3-z+AA). Stars mark the configurations carried forward as ApexFold-ESM and ApexFold-AF. **c,** Test accuracy as a function of the ESM-2 layer from which residue embeddings are extracted (L0, L12, L24, L30); intermediate-to-late layers (L12–L24) carry the most secondary-structure-relevant signal. **d,e,** Effect of ESM-2 backbone size (8M, 35M, 150M parameters) on accuracy **(d)** and MAE **(e)**. **f,** Effect of low-rank-adaptation (LoRA) rank (4, 8, 16) on accuracy across the `test`, Temporal-1 and Temporal-2 panels; the star marks the selected operating point. **g,** Test accuracy for two ESM-2 backbones (ESM-30, ESM-33) trained with and without loss normalisation. Here “Normalized” denotes mean-aggregating the reconstruction (task) loss *L*_recon_ over observed cells while leaving the auxiliary plasticity loss *L*_plast_ unchanged; normalisation improved the smaller backbone but not the larger one, so the unnormalized loss was used as the default. **h,** Accuracy when the AF3 pair (*z*) representation is progressively masked (full matrix, first half, second half, random 50%, random 25%), probing how much of the pair-track signal is required. **i,** Fraction of AF3 pair-representation energy contained within a diagonal band of increasing bandwidth (bw = 1, 3, 5), showing that the pair track is dominated by short-range, near-diagonal interactions consistent with local secondary-structure geometry.

